# Computational Study of Antibody Binding to SARS-CoV-2 Variants

**DOI:** 10.64898/2026.03.03.709420

**Authors:** Carolyn Chiu, M. Zaki Jawaid, Daniel L. Cox

**Affiliations:** Department of Physics and Astronomy, University of California, Davis, California; Epicrispr Biotechnologies, South San Francisco, California

**Author notes:** Department of Physics and Astronomy, University of California, Davis, California 95616.

**Keywords:** SAS-CoV2, antibody binding, immunity escape, simulation

## Abstract

**Background/Objectives:** The unprecedented structural and binding data for antibodies to the SARS-COV2 virus taken together with the mutations for the spike protein allows for a broad simulation study of antibody-spike protein binding. This provides an understanding of the co-evolution of human immunity and viral immunity escape.

**Methods:** We utilized the YASARA molecular dynamics program to generate initial antibody-spike structures and simulate to equilibration for six SARS-COV2 variants and 10 different antibodies sampling two different binding regions to the receptor binding domain of the spike (especially for the Class I antibodies in the same part of the spike which attaches to the ACE2 receptor protein) and one to the N-terminal of the spike. Starting structures for antibody binding to variant spike proteins are perturbatively achieved through point mutations and insertions/deletions in the YASARA program. We employed YASARA to measure interfacial hydrogen bound counts between antibodies and variant spike proteins, and the HawkDock MMGBSA program to characterize trends in binding energies with mutation for four of the antibodies. We utilized the VMD program to analyze the time course of hydrogen bond populations.

**Results:** As seen in previous studies, interfacial hydrogen bond counts serve as an excellent proxy for binding energies without the large systematic error inherent in the latter. We find that there is generally a decline in antibody binding strength, as measured by interfacial hydrogen bond counts, with viral evolution, but that a modest re-entrance of binding strength is present for most antibodies studied. Generically, the antibody heavy chain binds more strongly to the spike protein, through for approximately half the antibodies the light chain binding strength converges to the heavy chain strength with viral evolution.

**Conclusions:** The key conclusion is that the identified re-entrant immunity, speculatively arising from a balancing of maintenance of ACE2-spike binding while escaping antibodies through mutation, allows for some maintenance and even strengthening of immunity for later viral strains from early infection or vaccination.

## 0. Introduction

The SARS-CoV2 virus engendered a worldwide pandemic from March 2020 to May 2023 which killed tens of millions worldwide. The success of the virus in escaping immune responses through mutations of the SARS-CoV2 spike protein, which bound both to human angiotensin converting enzyme 2 (ACE2) receptor on the surface of cells and to potentially neutralizing antibodies, contributed to the lethality of disease over time.

Accordingly, there has been considerable experimental and computational interest in understanding which mutations through new strains of SARS-CoV2 were key to antibody escape. Many of the computational studies were oriented to specific antibody-spike interactions for particular strains of the SARS-CoV2 virus.

As has been noted elsewhere[1], there is a difficult evolutionary dance the virus plays to escape immunity: most of the effective neutralizing antibodies attach in the same receptor binding domain to which the virus binds to ACE2. Accordingly, a massive immune escape is likely to lead to a virus with weaker cell binding, and presumably lower cell lethality.

We have adopted a different perspective in this study, which computationally examines the binding strength of ten antibodies covering a broad range of binding sites to the spike protein, and examines six variants of the virus from the original strain of 2020 through the BA.2.86 variant predominant near the end of the pandemic in 2023. The antibodies studied include Class I (binding to the same set of residues in the spike receptor binding domain [RBD] as the ACE2), Class III (binding to the RBD but away from the ACE2 binding domain), and spike N-terminus binding. We are unaware of any study in the literature which has covered such a comprehensive range of antibodies and variants.

By monitoring the number of interfacial hydrogen bonds between the antibody and the spike protein, we are able to discern some new results relevant to the understanding of the pandemic trajectory.

First, while some antibodies display a monotonic decrease in binding strength with strain evolution, many show a partial reentrance, that is, the binding strength rebounds at least partially with time so that the antibodies retain some neutralizing capability. We believe this reflects the difficult evolutionary competition between immune escape and maintaining sufficient ACE2 binding. Second, quite uniformly the heavy chains of the antibody bind more strongly than the light chains. Third, in general, the Class I antibodies bind more strongly than the Class III or N-terminal antibodies studied. Fourth, those Class I antibodies alleged to show higher efficacy for omicron and descendant strains were not found to bind more strongly than the earlier delta or original (wild-type [WT]) strains.

Additionally, as we found in previous studies, the interfacial hydrogen bond count serves as a strong proxy for binding free energy which we evaluated separately for a representative subset of the antibodies.

The most important emerging qualitative picture from our study is that the viral evolution may provide immune escape from the current extant antibodies, but a global escape from all previous antibodies is likely impossible given that the most efficacious ones bind in the same region as the ACE2, so that high immune escape means weak cellular binding. Also because immune escape is relative to current extant antibodies, reentrance in which at there is at least some restoration of immunity from previous antibodies can lead to a persistent robust population immunity to the evolving virus.

## 1. Materials and Methods

### 1.1. Molecular Models

A summary of all the mutations relative to the original (WT) strain in the RBD and N-terminus of the spike protein from the six variants (Delta, BA.1, BA.2, XBB.15, BA.2.86) is found in Table 1.

**Table 1.** Mutations in variants relative to original (WT) strain including the D614G mutation. Column 1 lists the variant, Column 2 mutations in the receptor binding domain (RBD) and Column 3 mutations in the N-terminus. Mutations in the ACE2 binding region of Column 2 are identified in bold print. Point residue mutations were represented by XNY, where X is the WT residue, N the sequence number in the WT, and Y the variant residue. Deletions were represented by Δ(N-M), where *N* is the starting sequence number, *M* is the ending sequence number, and additions were represented by Σ(*N* − *M*) similarly. Sequences are listed at Ref. [2], with the original spike sequence at Ref. [3]

| Variant | RBD Mutations | N-Terminus Mutations |
| --- | --- | --- |
| Delta (B.1.617.2) | **L452R,T478K** | T19R,T95I,G142D,Y145H,<br>$\Delta(156-157)$,F158G,A222V,<br>W258L |
| BA.1 | G339D,S371L,S373P,S375F,<br>**K417N,N440K,G446S,S477N,**<br>**T478K,E484A,Q493R,G496S,**<br>**Q498R,N501Y,Y505H** | A67V,$\Delta(69,70)$,T95I,G142D,<br>$\Delta(143-145)$,N211K,$\Delta(212)$,<br>$\Sigma(R214)$ |
| BA.2 | G339D,S371F,S373P,S375F,<br>T376A,**D405N,R408S,K417N,**<br>**N440K,S477N,T478K,E484A,**<br>**Q493R,G496S,Q498R,N501Y,**<br>**Y505H** | T19I,L24S,$\Delta(25-27)$,G142D,<br>V213G |
| XBB.15 | G339H,R346T,L368I,S371F,<br>S373P,S375F,T376A,**D405N,**<br>**R408S,K417N,N440K,V445P,**<br>**E484A,G446S,N460K,S477N,**<br>**T478K,F486P,F490S,Q498R,**<br>**N501Y,Y505H** | T19I,L24S,$\Delta(25-27)$,G142D,<br>$\Delta(144)$,H146Q,Q183E,V213E,<br>G252V |
| BA.2.86 | G339H,K356T,S371F,S373P,<br>S375F,T376A,**R403K,D405N,**<br>**R408S,K417N,N440K,V445H,**<br>**G446S,N450D,L452W,N460K,**<br>**S477N,T488K,N481K,$\Delta(483)$,**<br>**E484K,F486P,Q498R,N501Y,**<br>**Y505H** | T19I,R21T,L24S,$\Delta(25-27)$,<br>S50L,$\Delta(69,70)$,V127F,G142D,<br>$\Delta(144)$,F157S,R158G,N211I,<br>$\Delta(212)$,V213G,L216F,H245N,<br>A264D,I332V |
| BA.2.75 | G339H,S371F,S373P,S375F,<br>T376A,**D405N,R408S,K417N,**<br>**N440K,G446S,N460K,S477N,**<br>**T478K,E494A,Q498R,N501Y** | T19I,L24S,$\Delta(25-27)$,G142D,<br>K147E,W152R,F175L,I210V,<br>V213G,G257S |

We drew starting structures for RBD-ACE2 binding from the Protein Data Bank. Class I antibodies bind in the same region of the RBD as the ACE2. Here we preferentially use the antibody designations in the literature, but we also refer parenthetically to the Protein Data Bank (PDB) files where the bound structures to relevant RBDs were displayed, P4A1 (7CJF)[4], C1A-B12 (7KFV)[5], 2-15((7L5B)[6], C1A-C2((7KFX)[5], C1A-F10(7KFY)[5], C1A-B3(7KFV)[5], S2X234(8ERQ)[7], and Omi3(7KF3)[8] were selected to represent the spectrum of Class I antibodies. Class III antibodies bind to the RBD away from where the ACE2 binds and is represented by CR.3022(6YOR)[9]. For antibodies that bind to the N-terminal domain, we used 4A8(7C2L)[10]. Fig. 1 shows the structures of representative spike domain-antibody complex types studied in this paper. The chosen antibodies were not comprehensive of all the known neutralizing antibodies for the SARS-CoV-2 spike but summarize a variety of antibodies that target the SARS-CoV-2 virus. We did not study T-cell binding. Antibodies were summarized in Table 2.

**Table 2.**
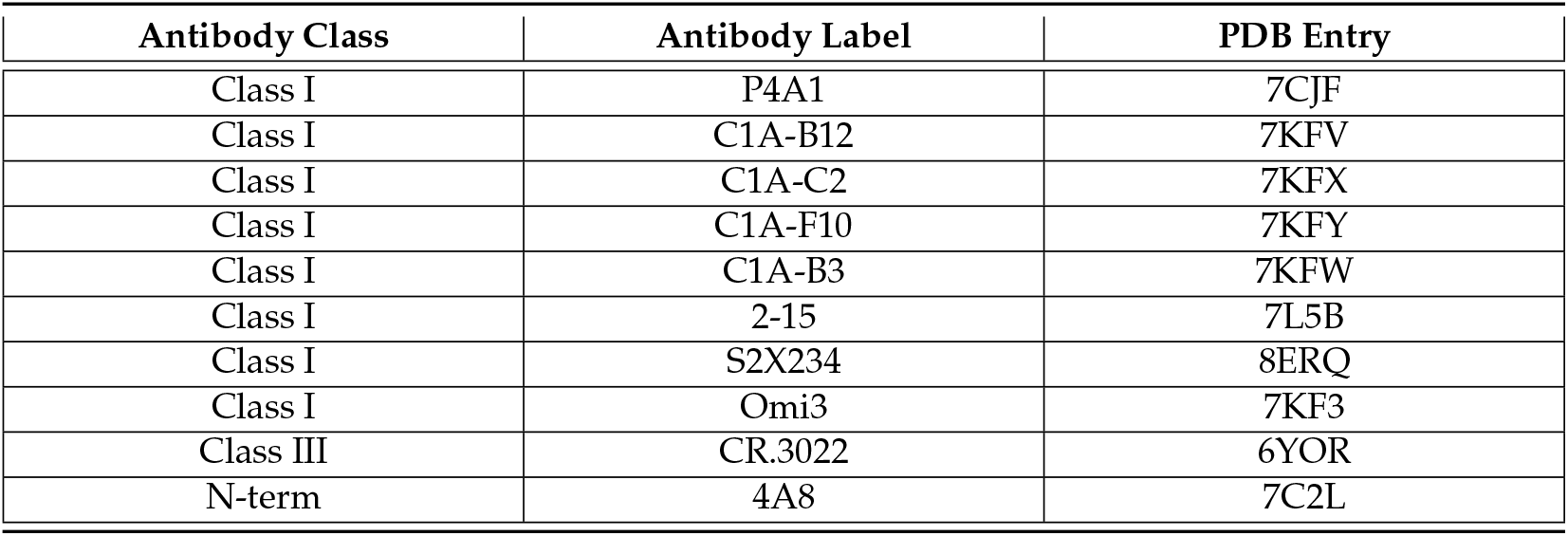
Antibodies by class studied here, with class in column 1, antibody nomenclature in column 2, relevant PDB entry in column 3.

**Figure 1.**
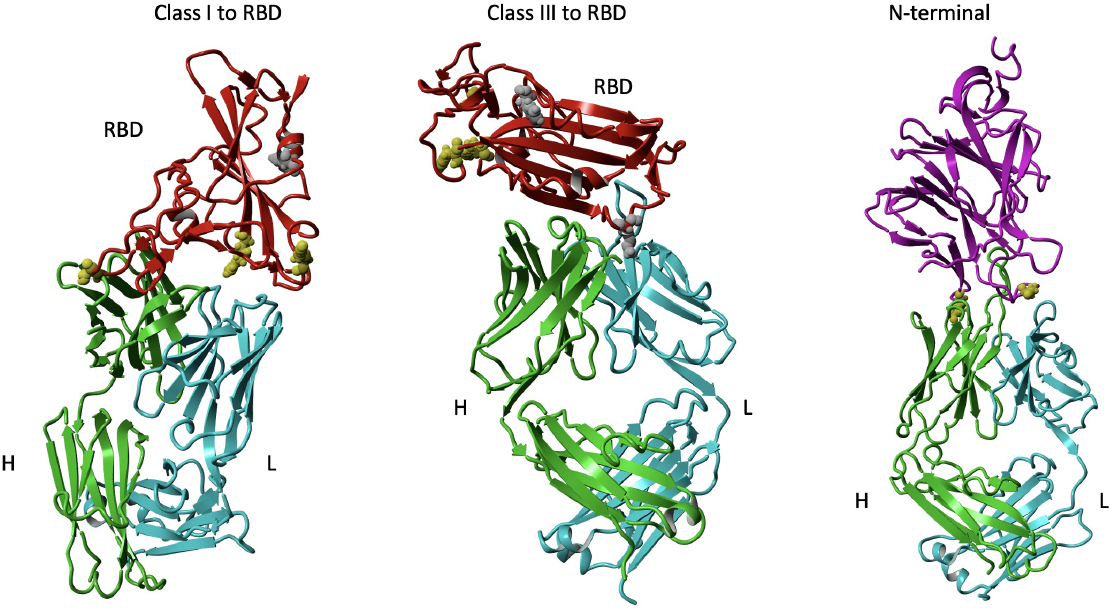
Structures of WT spike protein complexes studied. Binding of RBD (red) to Class I Ab P4A1 (binds in ACE2 interface region) and Class III Ab CR3022 (binds away from ACE2). For Class I, III binding, residue away from ACE2 (F347) shown in gray, residues in ACE2 binding region (R403,S477,Y505) shown in yellow. For CR3022 we also highlight K386 in gray Binding of NTD (purple) to Ab 4A8. Ab heavy chain green, light chain cyan. For N-terminal binding, residues near interface (K147,D253) shown in yellow.’ Graphic representations were created with YASARA[11]

To adopt starting model structures for studying binding with molecular dynamics, we picked the relevant earliest bound variant spike-antibody structure from the PDB and mutated the residues point by point within the YASARA modeling suite[11]. hree of these structures are shown in Fig. 1. When deletions arose, particularly in the N-terminus of the spike, we grafted the corresponding ends within YASARA[11]. Insertions and mutations were built upon starting structures using YASARA’s BuildLoop and SwapRes commands, respectively. While this is basically a perturbative approach biased to the starting structures, unlike an unbiased docking approach, it is unlikely to lead to systematic errors of the kind known in docking protocols.

### 1.2. Molecular Dynamics

Simulations of the protein-protein interactions were completed with the molecular-modeling package YASARA[11] by searching for minimum-energy conformations of the SARS-CoV-2-Ab complexes. For each structure, we carried out a energy minimization (EM) routine, which includes steepest descent and simulated annealing minimization to remove clashes and stabilize starting energies to within 50 J/mole.

All molecular-dynamics simulations were run using the AMBER14 force field with [12] for solute, GAFF2 [13], AM1BCC [14] for ligands, and TIP3P for water. The cutoff was 8 Å for Van der Waals forces (AMBER’s default value [15]) and no cutoff was applied for electrostatic forces (using the Particle Mesh Ewald algorithm [16]). The equations of motion were integrated with a multiple timestep of 1.25 fs for bonded interactions and 2.5 fs for non-bonded interactions at *T* = 298 K and *P* = 1 atm (NPT ensemble) via algorithms described in [17]. Prior to counting the hydrogen bonds and calculating the free energy, we carry out several pre-processing steps on the structure including an optimization of the hydrogen-bonding network [18] to increase the solute stability and a *pK*_a_ prediction to fine-tune the protonation states of protein residues at the chosen pH of 7.4 [17]. Simulation data is collected every 100 ps after at least 2 ns of equilibrium time, observed via the stabilization of: the number of hydrogen bonds, the root mean square deviations (RMSDs), and the interfacial surface area. For all simulations we require approximately 10 ns or more of equilibrated time as observed by stable values of root mean square deviation (RMSD) from the starting structure.

The total hydrogen bond (HBond) counts were tabulated using a distance and angle approximation between donor and acceptor atoms as described in [18] and averaged over the equilibration time series of the simulation. Results are shown in Fig. 2

**Figure 2.**
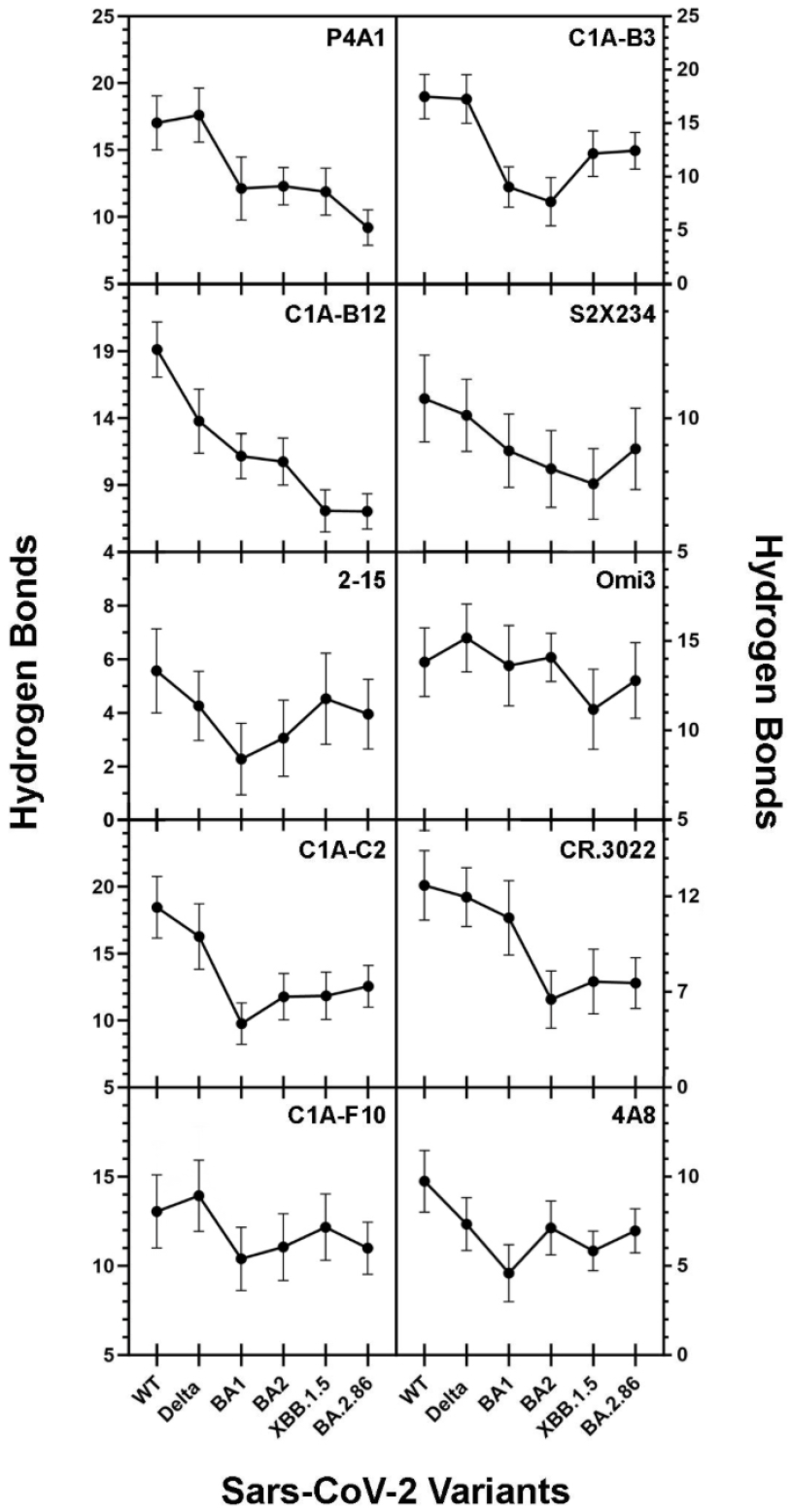
SARS CoV-2-Ab interfacial hydrogen bond counts. SARS Cov-2 variants WT, Delta, BA1 (omicron), BA2, XBB15, and BA.2.86 plotted for each Ab. BA.2.75 represents BA2 in 4A8 Ab graph. Note the reemergence effect for most variants where the binding strength rises after falling for subsequent variants. Graphing was performed with was performed using GraphPad Prism version 10.0.0 for Windows, GraphPad Software, Boston, Massachusetts USA, www.graphpad.com

Different equilibrium runs were generated by changing the starting random number seed within YASARA[11].

### 1.3. Endpoint Free Energy Analysis

Binding free energy for the energy-minimized structures from molecular dynamics simulations were calculated with the generalized Born surface area (MM/GBSA) method on the HawkDock server[19]. For Type I, Type III, and NTD antibodies, we average five snapshots of equilibrium conformations for binding to each SARS Cov-2 variant. The MM/GBSA approximations overestimate the magnitude of binding free energy in comparison to in-vitro experimental estimates, but correlate strongly with hydrogen bond counts. Correlation plots for endpoint free energy analysis against interfacial hydrogen bond counts are displayed in Fig. 3

**Figure 3.**
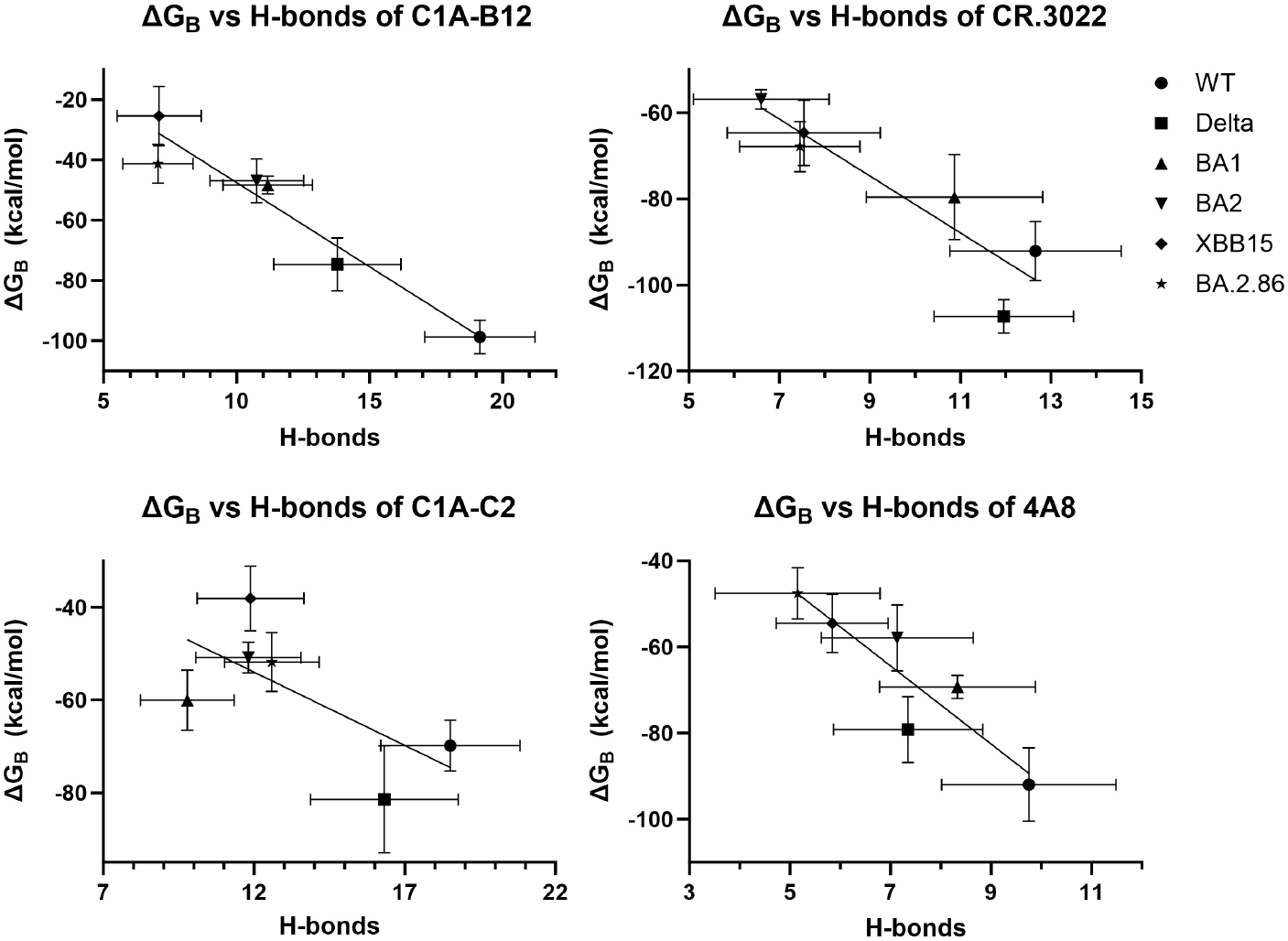
Binding free energy estimate in kcal/mole from GBSA analysis of molecular dynamics equilibrium conformations. Type I, Type II, and N-terminal Ab represented. Straight lines were from linear regression, with coefficients of determination for P4A1 *R*^2^ = 0.8713, C1A-C2 *R*^2^= 0.3731, CR.3022 *R*^2^= 0.7525, 4A8 *R*^2^ = 0.7056. Graphing was performed using GraphPad Prism version 10.0.0 for Windows, GraphPad Software, Boston, Massachusetts USA, www.graphpad.com

### 1.4. Statistical Significance

T-tests were performed on every combination of two hydrogen bond means with the same antibody and different spike protein variant. 15 combinations were compared for each antibody. For a given combination, their means, standard deviations, and number of points were input into the online T-test calculator by GraphPad[20] with the unpaired T-test selection. The difference in hydrogen bond counts were statistically significant when the generated p-value is less than 0.05. The p-value represents the probability that any difference between the two observed groups is due to random chance. A spread sheet of the t-test results are incluced in the Supplemental Materials.

### 1.5. Interfacial hydrogen bonds population analysis

YASARA[11] is effective at counting hydrogen bonds overall, which is a correlate to binding energy. To analyze the population of individual interfacial hydrogen bonds over the course of simulations, we employed a different strategy. First, we transformed the simulation snapshots to a GROMACS[21] file using the mdconvert macro[11]. Second, we uploaded the GROMACS trajectory to the Visual Molecular Dynamics (VMD) viewer[22].

VMD hydrogen bond analyses criteria differ in detail from YASARA. To provide the best match between the two separate programs we did the following. In the VMD menu, we chose the hydrogen bond analysis macro, with a fixed donor-hydrogen-acceptor (*D* −*H* −*A*) angle cutoff of *θ*_*c*_ = 35° (*θ*_*c*_ is actually 180° - ⟨*D* −*H* −*A*). YASARA softly cuts off for D-H-C anything for *θ*_*c*_ *<* 80°. YASARA imposes a distance criterion dependent upon the H-A distance[18], while VMD measures the D-A distance. Accordingly, we begin with a D-A default distance of 3.5 and vary the distance so that the average hydrogen bond count over the equilibrium trajectory matches that determined for YASARA. A full table of resultant hydrogen bond occupancies is available in online Supplemental Materials.

## 2. Results

### 2.1. Interfacial Hydrogen Bond Counts

The axis of variant is, effectively, an epidemiological timeline of COVID19 through the human population.

The most striking aspect of the interfacial hydrogen bond count vs variant for all but antibodies C1A-B12 and P4A1 is the non-monotonic “reentrant” behavior of the total hydrogen bond counts. Namely, after some initial decline from the earlier WT and Delta variants, there is subsequently some partial return in interfacial binding efficacy for later variants.

The results of t-tests show that this reentrance is statistically significant. We summarize as follows:

- **P4A1** Differences between BA1 and BA2 are not statistically significant (*p >* 0.05) nor are the differences between XBB15 and BA2.86. All other differences are statistically significant.
- **C1A-B3** Differences between WT and Delta, between BA1 and BA2, and between XBB15 and BA2.86 are not statistically significant, but all other differences are. Hence, the observed reentrance is statistically significant.
- **C1A-B12** Differences between BA1 and BA2, and differences between XBB15 and BA2.86 are not statistically significant, but all others are.
- **S2X234** Differences between WT and Delta are not statistically significant, but all other differences are, so that the observed reentrance for BA2.86 is statistically significant.
- **2-15** Differences between Delta and XBB15 are not statistically significant, but all other differences are, so the observed reentrance is statistically significant.
- **Omi3** Differences between WT and BA1, BA2 are not statistically significant, but all others are, sot he observed reentrance is statistically significant.
- **CA1-C2** Differences between BA2 and XBB15 are not statistically significant, all others are. Hence the observed reentrance for BA2, XBB15, and BA2.86 are statistically significant.
- **CR 3022** Differences between XBB15 and BA2.86 are not statistically significant, but all others are. Hence the reentrance for BA2.86 and XBB15 is statistically significant.
- **C1A-F10** Differences between BA2 and BA2.86 are not statistically significant but all others are. Hence the reentrance for BA2, XBB15, and BA2.86 is statistically significant.
- **4A8** Differences between Delta and BA2 are not statistically significant but all others are. Hence, the reentrance observed for BA2, XBB15, BA2.86 is statistically significant.

### 2.2. Binding Free Energy

We performed endpoint free energy analysis for the P4A1, C1A-C2, CR.3022, and 4A8 antibodies. As shown in Figure 3, with the exception of the C1A-C2 antibody, there is a high degree of correlation between the interfacial hydrogen bond counts and the endpoint free energy analysis. Hence, this continues the observation made in Refs. [1,23] that interfacial hydrogen bond counts are good proxies for endpoint free energy analyses.

The point is important because it can be seen that the binding free energies in Fig. 3 are quite large compared to values inferred from typical binding affinity data. As an example, we can take *K*_*D*_ ≈ 5 *nM* for ACE2-RBD binding from the literature[24]. The dissociation constant *K*_*D*_ is given by

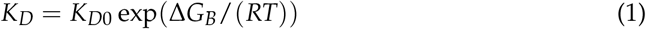

where standard estimates put *K*_*D*0_ ≈ 1 *M* (see, e.g., Dill and Bromberg[25]). Solving for Δ*G*_*B*_ gives −11.3 kcal/mole, clearly small in magnitude compared to the values found from GBSA analysis. Assuming the tighter binding of antibodies giving *K*_*D*_ = 0.1 *nM* changes the estimated Δ*G*_*B*_ to −14.9 kcal/mole, still far below the estimated magnitudes here.

The overestimates of the Δ*G*_*B*_ magnitudes derive from the GBSA approximation itself where large energies of opposite signs for the entire complex must cancel out to provide the binding energy. For example, for the CR 3022 WT binding, the GBSA contributions are, respectively: Van der Waals: −109.6 kcal/mole, Electrostatic: −302.7 kcal/mole, Generalized Born: 334.7 kcal/mole, and Surface Area: −14.6 kcal/mole. We anticipate the trends of the GBSA binding energy estimates to be accurate, but clearly the absolute values arising from the cancellation of opposing large energies are not.

### 2.3. Population Analysis of Hydrogen Bonds

Population analyses presented comprehensively in the Supplementary material, show that in every case, the majority of the interfacial hydrogen bonds are to the heavy chain of the antibody, although in a few cases, the bond strength to the light chain becomes comparable with viral evolution (for CR 3022, S2X234, C1A-B3, and 2-15 the light chain interfacial hydrogen bond count for the BA.286 is comparable to the heavy chain count).

There are no clear systematics about changing of hydrogen bonds with viral evolution. The predominant occupancies for each antibody and each variant are presented in the online supplemental materials.

## 3. Discussion

Unsurprisingly, this work shows a general diminution of binding by antibodies developed at a given time to the SARS-COV2 spike protein. Surprisingly, for many antibodies we have studied here there is a modest re-entrant behavior to the binding strength utilizing interfacial hydrogen bond count as a proxy per the correlation between the computed binding energy and the interfacial hydrogen bond count. Speculatively, this can be attributed to an volutionary drive to achieve antibody escape while maintaining reasonable binding of the receptor binding domain to the ACE2. However, the N-terminal antibody 4A8 and the Class III antibody CR 3022 show modest re-entrance This is subject to the caveat that our studies approached binding perturbatively from the original Wuhan strain of SARS-COV2 rather than entertaining a fully new binding motif. In all cases observed in simulation here, the reentrant behavior is statistically significant as measured by t-test p values.

This re-entrant immunity result is testable experimentally in affinity studies of antibody binding to SARS-COV2 variant spike proteins. When coupled with structural determinations from which interfacial hydrogen bond counts can be made an experimental test can be made of the correlation between binding free energy (log of the dissociation constant) and interfacial hydrogen bond count.

The significance of the re-entrant immunity is clear. Immunity gained from vaccination or earlier infection is not wholly surrendered, and newer antibodies developed to later variants or later vaccinations can maintain some efficacy against subsequent viral mutants. Meanwhile, since it is virtually impossible to strongly evolve away from Class I antibodies while maintaining sufficient ACE2 receptor binding to enter cells, there is general expectation for the potential damage by the virus to diminish with time, as discussed in previous work[1].

## 4. Acknowledgments

We acknowledge useful conversations with Rick Davis, Victor Muñoz, Javier Arsuaga, and Mariel Vazquez. Aspen Drake and Rustin Mahboubi-Ardakani contributed to some of the simulations.

We thank J. Solana, M. Vazquez, and R.L. Davis for useful discussions in early phases of this work. A. Drake and R. Mahboubi-Ardakani contributed to early work on the simulations.

## Author Contributions

The study design was initiated by M.Z. Jawaid and D.L. Cox, and later work by C. Chiu. Simulations were carried out by C. Chiu (60%), D.L. Cox (30%), and M.Z. Jawaid (10%). Statistical analyses and endpoint free energy analyses were performed by C. Chiu. H-bond population analyses were performed by D.L. Cox. The manuscript was written by C. Chiu and D.L. Cox, with editing by M.Z. Jawaid.

## Data Availability Statement

Data for simulations, analyses, and hydrogen bond populations is available at https://drive.google.com/drive/folders/1ZZg0VOnmag9k8El5af459iiHAzPapdz?usp = *sharing*

